# Variation of Myelin-associated Gene Expression and *Ndrg1* within the Prefrontal Cortex as Determinants for Initial Level of Response to Alcohol

**DOI:** 10.1101/2025.10.19.683339

**Authors:** Jennifer T. Wolstenholme, Sean P. Farris, So Hyun Park, Andrew D. Van der Vaart, Michael F. Miles

## Abstract

**Background:** Studies in humans and animal models have documented relationships between initial sensitivity to alcohol and alcohol drinking behavior.

Prior expression profiling studies of C57BL/6J and DBA/2J mice and rhesus macaques within the prefrontal cortex (PFC) have shown variation in myelin gene expression may be linked with alcohol sensitivity and consumption.

**Methods:** Combining gene expression studies from human and mouse PFC, we identified a cross-species gene network enriched for myelin-associated genes. Since myelin expression is correlated to alcohol sensitivity and alcohol drinking behavior, we hypothesized basal levels of PFC myelin gene expression may be a genomic determinant for these behavioral responses. Using an animal model of CNS demyelination, and localized knock down of N-myc downstream regulated gene 1 (*Ndrg1)*, we measured effects of cortical myelin reduction on initial alcohol sensitivity and drinking behavior.

**Results:** Reducing myelin-related gene expression significantly altered sensitivity to alcohol and decreased alcohol consumption. Mouse genetic-based studies identified *Ndrg1* as a putative quantitative trait gene for sedative-hypnotic responses to alcohol. Site-specific injections of shNdrg1 lentivirus into PFC led to a significant decrease in NDRG1 expression, causing increased alcohol behavioral sensitivity and reduced preference for high concentrations of alcohol.

**Conclusion:** Myelin is an important biological component underlying CNS disorders. Our studies demonstrate the role of a novel candidate gene (*Ndrg1*), and myelin-associated gene expression, as an important factor modulating initial sensitivity to alcohol and alcohol consumption. Differences in the expression of myelin-related genes, including *Ndrg1*, may serve as future therapeutic targets for the treatment of alcohol use disorders.

## Introduction

Molecular programming and functional circuitry of the central nervous system (CNS) influences the development of psychiatric and neurological disorders. Defining coordinate biological perturbations occurring within defined regions of the CNS is important for understanding the etiology of disease. An alcohol use disorder (AUD), similar to other psychiatric disorders, is driven by multiple interacting genetic systems, molecular networks, and intermediate behavioral phentoypes. Human studies and animals models for AUDs have previously shown the risk of excessive alcohol consumption is related to the initial level of response to alcohol (1; 2). Systematic identification of mutual processes affecting initial alcohol sensitivity and consumption is essential for helping mitigating disease burden of an AUD.

CNS studies of the prefrontal cortex (PFC) have shown coordinate regulation of myelin-associated gene expression by ethanol in humans (3–5) and animal models (6). Neuroimaging studies have confirmed the role of CNS myelin in AUD, strongly suggesting alcohol consumption is correlated with myelin structure in frontal cortex (7). Primate studies by our laboratory showed robust increases in connectivity of a myelin gene rich expression network in animals consuming chronic ethanol and implicated N-myc downregulated gene-1 (*Ndrg1*) as a major hub gene in such animals. NDRG1 is a myelin-associated protein and mutations in the coding region of this gene produce a demyelinating polyneuropathy. Dysfunction of myelin in discrete brain regions also confers specific cognitive and motor impairments (8). Prospective neuroimaging studies, begun prior to any alcohol drinking in adolescents progressing to heavy drinking, suggests myelin and alcohol consumption are interlinked (9; 10) (11). Impairments in myelin expression and formation likely disrupt axonal integrity, putting individuals at risk for substance abuse (12; 13) as well as cognitive impairments (14; 15).

Myelin dysregulation has also been implicated in many other neuropsychiatric disorders, including schizophrenia (16; 17), major depression (18), and cocaine abuse (19–21). Dysregulation of myelin-associated gene expression in multiple disorders may indicate a common molecular substrate in the neurobiology of disease (22). The functional relationship between neuropsychiatric disorders and CNS myelin is unknown, although it likely affects both white matter and neuronal integrity (23). Myelin is a dynamic process, meeting cellular requirements to achieve efficient CNS communication for interacting with the environment. Dysfunction of myelin-related molecular networks creates a pathogenic loop affecting neuronal signaling and neuropsychiatric disorders, including alcohol abuse and dependence. However, it is unclear if disruptions in myelin abundance are a causal predisposing factor for initial alcohol sensitivity and consumption. Differences in alcohol-responsive phenotypes may be influenced by baseline variation in myelin-related gene expression, which may contribute to the pathogenesis of an AUD.

Herein we demonstrate variation in the abundance of myelin-associated gene expression within PFC may be predictive indicator of acute alcohol sensitivity and consumption. Although it is difficult to establish a causative association between variation in gene expression and complex traits, convergent evidence from multiple models can support causal relationships (24; 25). Cross-species analysis facilitates a comprehensive translational approach for the prioritization of novel candidate genes and gene networks underlying complex traits (26; 27). Our results support the hypothesis that a myelin-associated gene network involving *Ndrg1* within PFC is an important factor modulating CNS plasticity related to initial alcohol sensitivity and alcohol drinking behavior.

## Methods and Materials

### Gene expression data analysis

#### Differential gene expression analysis

All animals were treated according to protocols for animal care established by Virginia Commonwealth University and the National Institute of Health. Total RNA isolated from PFC was analyzed using whole-genome Affymetrix microarrays (n=3/group). Brain dissection and RNA isolation was performed as previously described (6). Significance-scores (S-scores) were computed for comparing gene expression differences between: ISS vs ISS, B6 vs D2, ISS vs S.5L, and ILS vs L.5S. One-class statistical analysis of microarrays was applied to determine differential gene expression that is significantly different from 0 using a 5% false-discovering rate (FDR) (28). Baseline differences were further filtered for a mean S-score ≥ 1.5 or ≤ -1.5 (composite significance of *P* < 0.01). Gene enrichment analysis of gene lists was conducted using the ToppGene Suite (29).

#### Gene expression analysis of BXD and LXS RI mice

PFC gene microarray expression data has been previously generated by our laboratory for adult male BXD RI (n=29) and LXS RI (n=42) PFC and by University of Colorado for RNAseq data on LXS RI, ILS and ISS whole brain (n=44 strains). Gene expression data for BXD PFC (GN135), LXS PFC (GN130), LXS hippocampus (GN133) and LXS whole brain (GN783) are deposited in the GeneNetwork resource (http://genenetwork.org/webqtl/main.py) (30). GeneNetwork also contains a database of recorded phenotypes for BXD and LXS RI mice. Each phenotype has been assigned an identifier, corresponding to a respective published report (if available). Behavioral phenotypes (n=4,372 for BXD and n=189 for LXS mice) for both RI lines were correlated with all PFC RMA intensity values (31) that possessed a sufficient number of common mouse strains. Duplicate gene symbols were removed to retain gene expression measurements with the largest statistical significance. Genetical-genomic analysis of *Ndrg1* across the LXS RI panel was done using RMA intensity scores to generate an interval map for *Ndrg1,* with significance levels determined by permutation analysis. LXS whole-brain (n=62) and hippocampus (GN133, n=79) gene expression was used for additional cis-eQTL analysis. LXS whole-brain data was obtained from Phenogen (https://phenogen.ucdenver.edu/PhenoGen/) (32) and is also deposited in GeneNetwork as above.

#### Human gene expression

PFC gene expression was retrieved from Gene Expression Omnibus (GEO: GSE29555) for alcoholic and non-alcoholic brain tissue (33; 34). Expression profiling was performed using the Illumina HumanHT-12 V3.0 Expression BeadChip from fresh-frozen sections of tissue from the superior prefrontal cortex. Cases have been previously matched as closely as possible by age, gender, postmortem interval, brain pH and smoking history.

### Reduction of myelin by cuprizone

Adult male C57BL/6J mice (6 to 7 week-old, Jackson Laboratory, Bar Harbor, ME) were singly housed on a 12-hour light/dark cycle with *ad libitum* access to water and standard rodent chow (catalog #7912; Harlan Teklad, Madison, WI) or 0.2% cuprizone (w/w, Bis(cyclohexanone)oxaldihydrazone; Sigma) added to standard rodent chow. Mice were fed +/- cuprizone for five weeks to induce mild-to-moderate demyelination (35), and were maintained on their respective diet throughout behavioral testing. Separate groups were tested for myelin-related gene expression, immunohistochemistry, and behavioral measures. Mice were tested for locomotor activity, alcohol drinking behavior, alcohol-induced loss of righting (LORR), handling induced convulsions, and gait analysis (see Supplemental Methods).

### *In vivo* knockdown of *Ndrg1*

Adult male C57BL/6J mice were infected with shNdrg1 or scrambled virus (pLL3.7-SC) by stereotaxic surgery (36). Ndrg1-kd lentivirus or pLL3.7-SC 7x 10^-6^ TU/μl (Supplemental Methods) was injected into PFC (0.6mm lateral, 1mm anterior, and 2mm ventral to bregma). A subset of mice were sacrificed 3-weeks following surgery to verify the site of injection, eGFP expression, and NDRG1 protein knockdown via immunohistochemistry. Remaining mice were tested for chronic alcohol-drinking behavior via an intermittent access three-bottle-choice paradigm, and alcohol-induced LORR (see Supplemental Methods).

### Statistical Analysis

All data are reported as mean ± standard error and analyzed using Student’s t-test, One-way, Two-way ANOVA or Repeated Measures ANOVA for each independent assessment. Post-hoc comparisons were conducted using Student Newman-Keul’s test. *P*-values ≤ 0.05 were considered statistically significant.

## Results

### Identification of basal differential gene expression within PFC

Gene expression studies from our laboratory and others have demonstrated variation across different model organisms and brain regions; which may influence neuropsychiatric phenotypes such as predisposition for alcohol consumption and related behaviors. C57BL/6J (B6) and DBA/2J (D2) mice demonstrate a number of differences in alcohol-responsive traits. Prior gene expression profiling of the mescorticolimbic dopaminergic reward pathway, spanning prefrontal cortex (PFC), nucleus accumbens (NAC), and ventral tegmental area (VTA) shows baseline differences in several clusters of coordinately regulated genes relevant to the neurobiology of addiction. To extend our previous studies we sought to characterize basal variation of gene expression involving a specific behavioral phenotype, alcohol-induced loss of righting reflex (LORR). LORR is a sensitive measure of an animals sleep time in response to an acute high-dose exposure of alcohol (37). Inbred short-sleep (ISS) and inbred long-sleep (ILS) mice have been selectively bred to help discern underlying determinants involved in acute alcohol sensitivity. Our analysis focused on PFC, a brain region encompassing the anterior cingulate and primary motor cortex that has a known involvement in the neurobiology of addiction and executive motor function.

Unbiased profiling of ISS and ILS mouse PFC determined 952 genes were significantly altered between these two genetically divergent strains. A significant number of genes (n=95, *P* = 3.19 e-35) were also differentially expressed at baseline between B6 and D2 mouse PFC (Fig. 1A; Supplementary Table 1), suggesting a common set of genes related to differences in alcohol behavioral sensitivity across four strains of mice (i.e. B6 vs D2 & ISS vs ILS). The overlapping gene set was enriched for members of the ‘myelin sheath’ (GO:0043209, *P* = 5.01 e-06) and ‘glutamate receptor signaling’ (GO:0007215, *P* = 2.75 e-05) (Supplementary Table 2). Genes involving glutamate receptors, and related signaling mechanisms, have been previously implicated in alcohol behavioral phenotypes, including LORR (38; 39). Additionally, our work on ethanol consumption in a primate model has previously shown that a expression network significantly enriched with myelin genes showed correlation with ethanol consumption and significant network connectivity changes between control and ethanol consuming animals. In particular, a gene previously identified as responding to acute ethanol and also contained in the group of 95 genes commonly differently expressed between ILS vs. ISS and B6 vs. D2 mice, Ndrg1, was found to be a major hub gene for the primate myelin network of ethanol consuming animals. Combined, these data suggest that Ndrg1 and myelin gene expression more generally, might be important molecular substrates influencing initial alcohol sensitivity and long-term behaviors such as chronic consumption.

**Figure 1.**
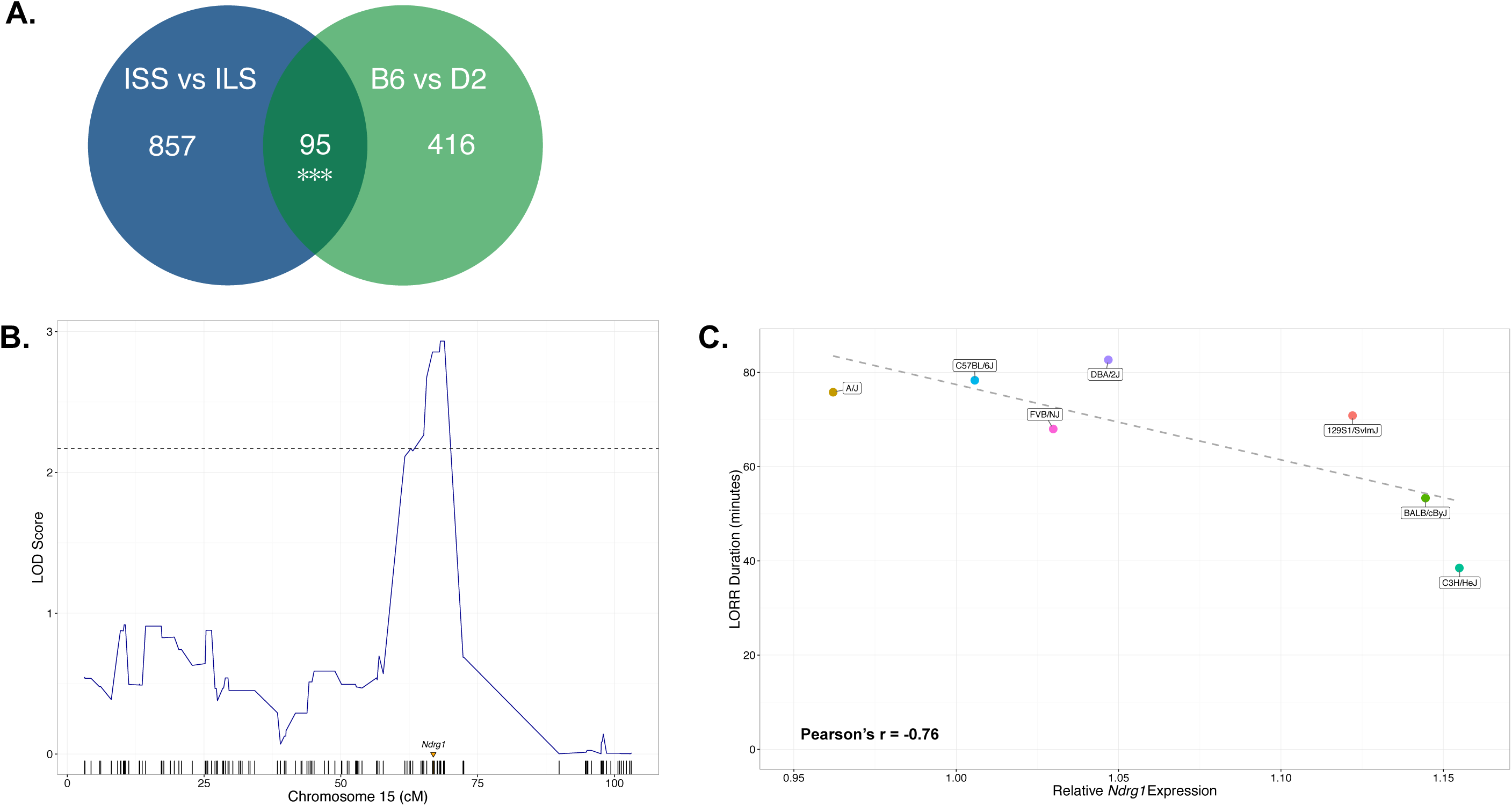
Prioritization of *Ndrg1* within the PFC for alcohol-induced LORR. **(A)** Venn-diagram showing the number of overlapping genes with basal differentially expressed genes for ISS versus ILS and for B6 versus D2 mouse PFC. Further analysis of basal differences within the PFC using reciprocal cogenics mice, ILS.ISS-*Lore5* (L.5S) and ISS.ILS-*Lore5* (S.5L), limiting our focus to differences in the expression of *Ndrg1*. **(B)** Variation in basal PFC expression of *Ndrg1* across LXS mice (n=42), showing a potential cis-eQTL for *Ndrg1* within this mouse population. Additional cis-eQTL analysis for *Ndrg1* across two independent brain studies of LXS mice is shown in Supplementary Fig. 1. **(C)** Variation in *Ndrg1* expression within the PFC is inversely correlated (Pearson’s r = -0.76, *P* = 4.7 e-02) with the mean alcohol-induced duration of LORR across A/J, C57BL/6J, FVB/NJ, DBA/2J, 129S1/SvlmJ, BALB/cByJ, and C3H/HeJ mice (n=5-6/strain).

We initially investigated *Ndrg1* differential expression between ILS and ISS mice by probing whether this difference was due to cis-acting genetic variants influencing **Ndrg1** expression. PFC expression of *Ndrg1* varied across the LXS recombinant inbred (RI) panel (n=42) in GeneNetwork data. Derived from ILS and ISS progenitors, LXS mice showed evidence of a cis quantitative trait locus for *Ndrg1* (Fig. 1B). A cis-eQTL for *Ndrg1* was replicated across two independent LXS RI CNS datasets (Supplementary Fig. 1A & 1B). *Ndrg1* is located within a non-identical by descent region of the ILS / ISS genome (Supplementary Fig. 1C); suggesting a nearby polymorphism may affect the regulation of *Ndrg1* expression within this selectively bred mouse population. Due in part to the mosaic genetic architecture existing among laboratory mice, a portion of nucleotide variants may be strain specific and not universally shared among differing populations (42; 43); however, biological variation of transcribed genes, serving as an important molecular intermediate, may affect behavioral phenotypes (44; 45). To test this hypothesis we examined variation in the expression of *Ndrg1* across seven commonly used inbred laboratory strains, previously shown to vary in their response to acute alcohol induced LORR (46). LORR was significantly correlated with the previous report (Supplementary Fig. 2; Pearson’s r = 0.91, *P* = 4.8 e-03), indicating variation of LORR among these mice is consistent in different environments. Basal expression of *Ndrg1* was inversely correlated with LORR duration (Fig. 1C; Pearson’s r = -0.76, *P* = 4.7 e-02). As determined by the LORR, variation in the baseline expression of *Ndrg1* may be an important molecular endophenotype influencing initial sensitivity to alcohol.

### Characterization of an *Ndrg1* gene coexpression network and alcohol-related behaviors

To provide a biological context for *Ndrg1* we constructed a gene coexpression network, wherein co-regulated transcripts impart molecular systems relevant to *Ndrg1* and alcohol-related phenotypes. The *Ndrg1* coexpression network was determined across LXS (n=42) and BXD (n=29) RI lines, which respectively originate from the parent strains used to initially prioritize *Ndrg1* (i.e. B6 x D2 and ILS x ISS). To limit pairwise gene-gene relationships that are most likely conserved across species and relevant to the human condition, *NDRG1* correlations were also determined from human PFC (n=31)(33). Anchored by *NDRG1*, intersecting significant correlations (p<0.05) across LXS, BXD, and human PFC datasets determined a coexpression network involving 1,485 genes (Supplementary Table 3).

To discern potential molecular underpinnings of alcohol-related behaviors we examined co-variation in PFC gene expression in relation to LXS and BXD mouse phenotypes (Supplementary Table 4). The *Ndrg1*-network was significantly enriched for PFC genes mutually correlated to multiple acute alcohol-induced LORR behavioral studies conducted in LXS and BXD mice (Fig. 2A). Additionally, the *Ndrg1*-network was enriched for PFC genes mutually correlated to animal models measuring alcohol intake (Fig. 2A). Although acute alcohol behavioral information is unavailable for postmortem human brain studies, using estimates of lifetime alcohol consumption we determined the *NDRG1*-network relationship with alcohol intake was conserved within human PFC (Fig. 2A). Consistent enrichment of the Ndrg1-network across mouse and human PFC could suggest an underlying molecular relationship with acute alcohol sensitivity and chronic alcohol consumption.

**Figure 2.**
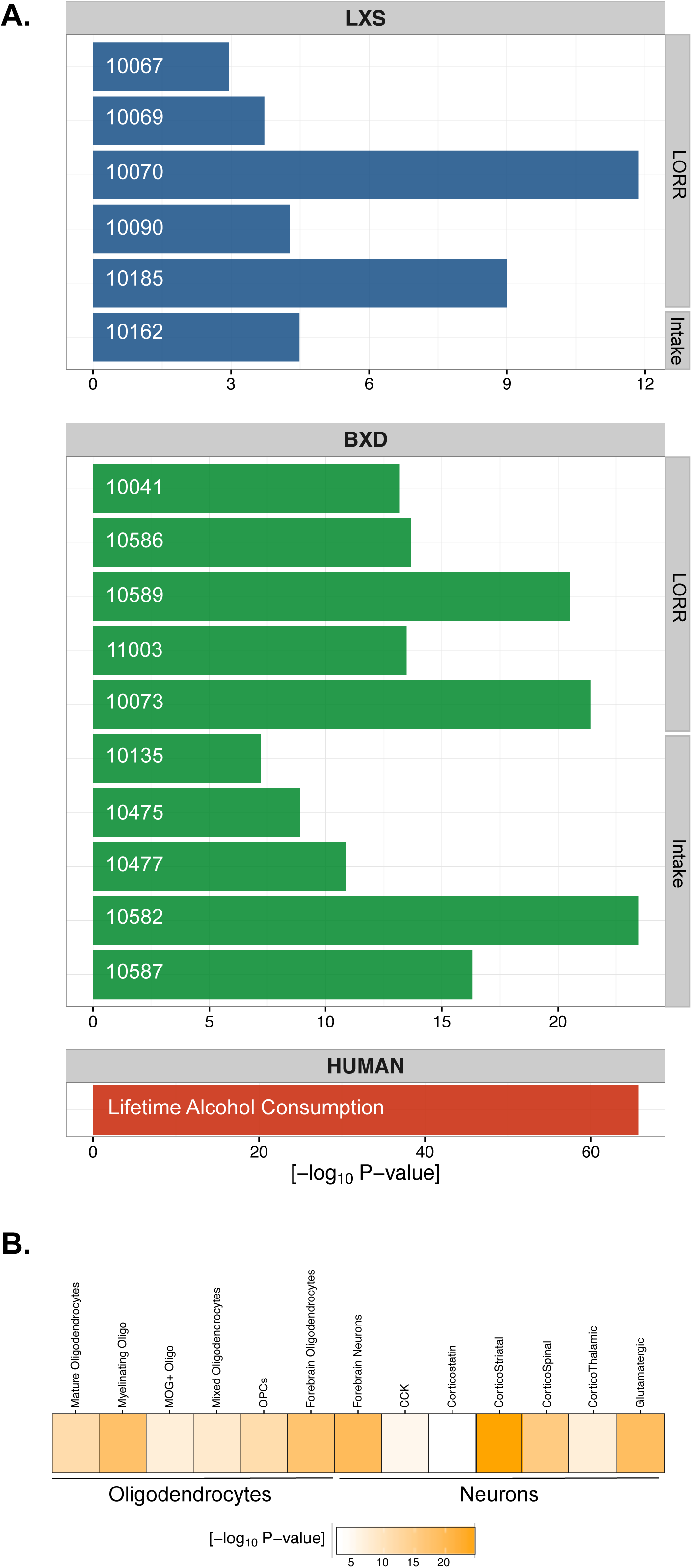
Identification and characterization of a conserved *Ndrg1* gene coexpression network within the PFC. **(A)** *Ndrg1* coexpression network of 1,485 genes within the PFC, with genes indicated as circles and correlations represented as lines between genes. **(B)** Over-representation of alcohol-related behavioral phenotypes for genes mutually correlated to *Ndrg1* within the PFC and available phenotypes. Data is shown for LXS mice (green), BXD mice (blue), and estimates for lifetime alcohol consumption in the human samples studied herein. Data are labeled using GeneNetwork Record IDs for phenotypes relevant to the LORR and alcohol intake are shown along the right-vertical axis, with full results reported in Supplementary Table 4. **(C)** Heatmap showing over-representation of the *Ndrg1* gene coexpression network for genes expressed in oligodendrocytes and neurons (82–84).

The molecular function of *Ndrg1* within the CNS is largely unknown; however, deficiencies or mutations in *Ndrg1* can produce a progressive demyelination phenotype (47; 48) [1, 2]. The *Ndrg1* coexpression network contains a significant number of genes within the ‘myelin sheath’ (GO:0043209, *P* = 6.23 e-19), further suggesting *Ndrg1* plays an important role in CNS myelination. Specialized glial cells in the CNS, known as oligodendrocytes, are responsible for the production of myelin that ensheaths neuronal axons to facilitate efficient neuronal communication through the propagation of action potentials. The *Ndrg1*-network contained a significant fraction of genes expressed within differing stages of oligodendroglia maturation (Fig. 2B), including oligodendrocytes precursor cells (OPCs) (*P =* 4.52 e-14), myelinating oligodendrocytes (*P =* 1.02 e-20), and mature oligodendrocytes (*P* = 3.01 e-14). Cell-type specific markers of oligodendroglia and major components of the myelin sheath (i.e. *Mobp, Mal, Mbp, Plp1, Mog, Mag*) are among the top genes consistently correlated with *Ndrg1* expression (Supplementary Table 3). Variation in expression of the *Ndrg1*-network may thus affect neuronal-glia processes involved in the production and maintenance of myelin, capable of influencing neuronal firing rates and CNS-mediated behaviors.

### Validation of myelin-related gene expression for LORR and alcohol consumption

Cross-species analysis of *Ndrg1* gene coexpression network identified a coordinately regulated network of myelin-associated genes in PFC, providing convergent evidence for basal variation of myelin-associated genes as an underlying factor in initial alcohol sensitivity and alcohol consumption. To validate the role of myelin in alcohol-related behaviors, C57BL/6J mice were placed on a 0.2% (w/w) cuprizone-supplemented diet to induce a demyelination phenotype. Cuprizone selectively targets oligodendrocytes, causing a predominantly frontal demyelination in adult B6 mice (49). Following 5-weeks of cuprizone, seven myelin-associated genes were significantly decreased in the PFC (Fig. 3A, *P* < 0.05 for *Ndrg1*, *Mobp*, *Mbp*, *Plp*, *Mag*, *Mog*, and *Mal*). *Ndrg1* expression co-varies with chloride intracellular channel 4 (*Clic4*) within PFC (Supplementary Table 3), but exposure to cuprizone did not alter *Clic4* (*P* = 2.87 e-01) (Fig. 3A). Selectively targeting neuronal expression of *Clic4* within PFC significantly changes the duration of alcohol-induced LORR (36); suggesting the effects of cuprizone were largely restricted to oligodendroglia. Immunostaining of coronal sections with the selective myelin stain fluoromyelin (50), demonstrated a decrease in width of the corpus callosum, and qualitative thinning of myelinated fibers extending into the PFC (Fig. 3B). Coordinate thinning of myelin, and decreased abundance of seven myelin-related genes in PFC indicates cuprizone was effective in reducing expression of the myelin-associated gene network. Importantly, previous studies have shown cuprizone predominantly causes a down-regulated myelin expression within PFC, without affecting other more posterior brain regions such as striatum (35).

**Figure 3:**
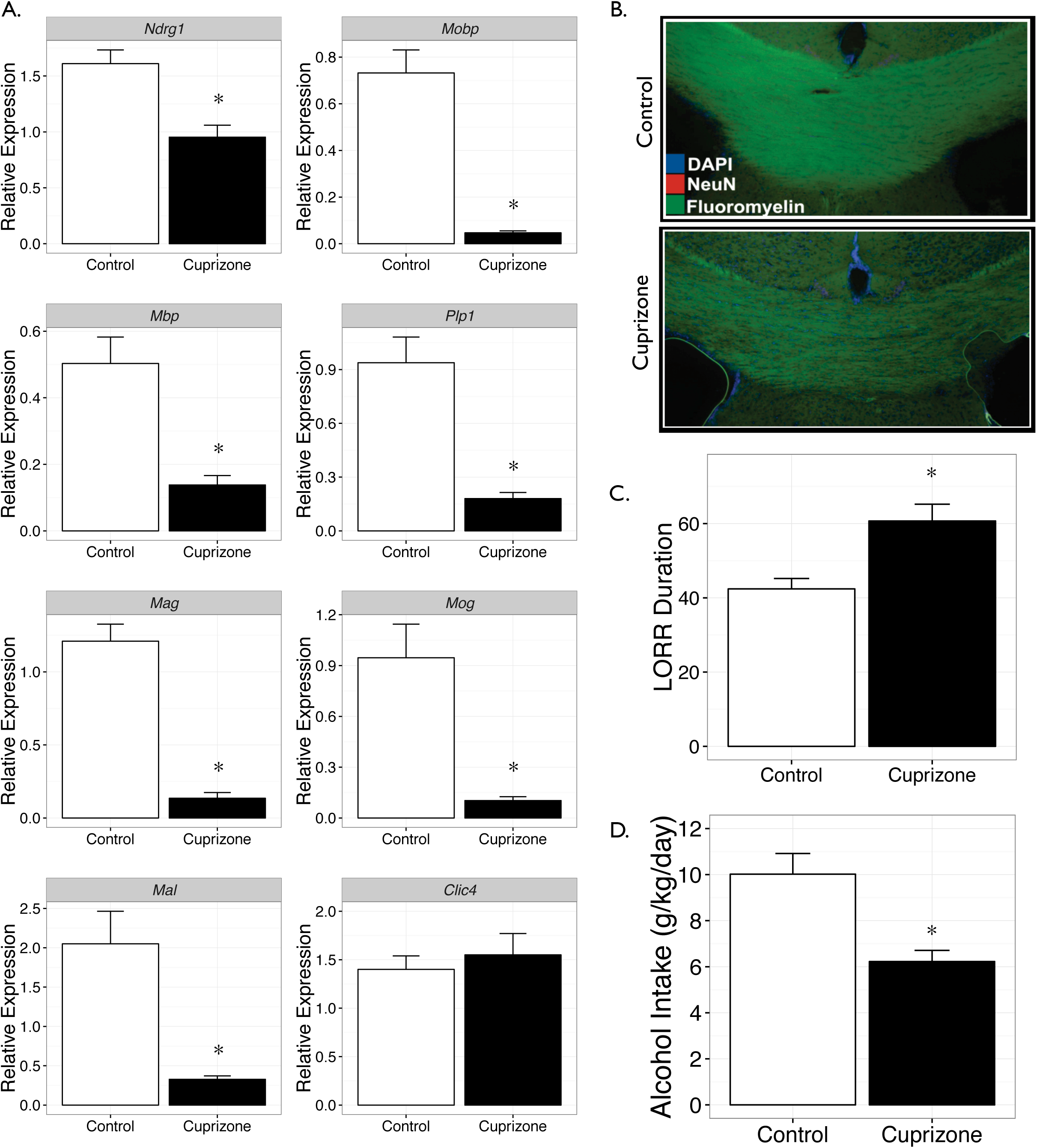
Cuprizone decreases myelin network genes and alters alcohol behavior. **(A)** Relative mRNA expression for *Ndrg1* (*P* = 1.6 e-02), *Mobp* (*P* < 1.0 e-04)*, Mbp* (*P* = 7.0 e-04)*, Plp1* (*P* = 2.0 e-04)*, Mag* (*P* < 1.0 e-04)*, Mog* (*P* = 9.0 e-04)*, Mal* (*P* = 1.0 e-03), and *Clic4* (*P* = 2.87 e-01), demonstrating a selective decrease for basal expression of myelin-associated gene expression in mouse PFC (n=7-8/group). **(B)** Representative immunostaining using Invitrogen BrainStain imaging kit for DAPI (blue), NeuN (red, no visible staining in these sections), and fluoromyelin (green) showing decreased myelin content in the corpus callosum of mice fed a cuprizone diet. **(C)** Cuprizone increased the mean duration of alcohol-induced LORR behavior compared to controls (*P* = 2.9 e-03, n=10/group). **(D)** Cuprizone decreased alcohol intake in a 2-bottle choice task (*P* = 1.5 e-03, n=15/group). Data shown as Mean ± SE. * *P* < 0.05 versus control.

In agreement with our gene expression profiling of the PFC across RI panels, cuprizone caused changes in select alcohol behaviors (*summarized in* Supplementary Table 5). Sensitivity to the sedative-hypnotic effects of acute alcohol was increased for cuprizone mice. Cuprizone-exposed mice displayed a significantly longer LORR duration compared to mice on a control diet (Fig. 3C. *P* = 2.9 e-03). Controls had a mean LORR duration equal to 42.4 minutes, compared to a mean LORR duration of 60.7 minutes in cuprizone mice. The onset for LORR was similar between groups (∼85 seconds, *P* = 5.56 e-01). Cuprizone mice consumed 36% less alcohol than controls in a 2-bottle choice test (Fig. 3D, *P* =1.5 e-03), and showed reduced alcohol preference (*P* < 1.0 e-04). In taste perception assays comparing saccharin and quinine versus water, cuprizone did not alter the preference for either tastant (Supplemental Fig 3, *P* =3.6 e-01 for saccharin and *P* = 3.23 e-01 for quinine). Alterations in LORR and alcohol drinking behavior cannot be explained by non-specific effects on alcohol metabolism because cuprizone did not alter the pharmacokinetics of blood ethanol concentrations (BEC) (Supplementary Fig. 4).

Behavioral differences were also not explained by confounding effects of cuprizone on motor function. There were no significant effects of cuprizone on locomotor-related or gaiting behaviors (Supplementary Fig. 5). Cuprizone also had no effect on generalized CNS excitability, showing no significant differences in baseline handling-induced convulsions, or withdrawal from acute alcohol exposure (Supplementary Fig. 6). Behavioral validation of this model for reduced CNS myelin demonstrates a selective effect for high-dose alcohol exposure (i.e. LORR) and voluntary alcohol consumption. These results connote a conserved coexpression network within the PFC involving multiple myelin-related genes (including *Ndrg1*) are an underlying component in initial alcohol sensitivity and alcohol consumption.

### Validation of the candidate gene *Ndrg1* for LORR and alcohol consumption

Genomic analysis across multiple inbred mouse strains identified variation in *Ndrg1* expression as a possible factor for differences in acute alcohol sensitivity. The magnitude of change in *Ndrg1* expression, although reduced, was less affected by cuprizone in comparison to other myelin-related genes examined (Fig. 2A); suggesting *Ndrg1* may be expressed in additional non-myelinating CNS cell-types. The *Ndrg1* coexpression network is also enriched within several neuronal-related subtypes (Fig. 2B). To selectively probe the individual role of *Ndrg1* in acute alcohol sensitivity and drinking behavior we developed a lentivirus shRNA vector (pLL3.7-Ndrg1-kd) to specifically decrease PFC *Ndrg1* expression within a broad spectrum of cell-types. Transformation using NIH/3T3 cells was used to confirm a significant reduction of *Ndrg1* mRNA and protein expression following transduction with pLL3.7-Ndrg1-kd (Supplementary Fig. 7). Compared to NIH/3T3 cells transduced with scrambled control (pLL3.7-SC), *Ndrg1* expression was reduced by 52% in cells transduced by pLL3.7-Ndrg1-kd (*P* = 1.5 e-02, Supplementary Fig. 7A). Protein expression was decreased 71% by pLL3.7-Ndrg-kd relative to pLL3.7-SC (*P* = 3.08 e-02, Supplementary Fig. 7B). Site-specific pLL3.7-Ndrg1-kd administration into B6 PFC showed *in vivo* reduction of NDRG1 protein expression by immunohistochemistry (Supplementary Fig. 7C).

Lentiviral injections into medial PFC of B6 mice with pLL3.7-*Ndrg1*-kd or pLL 3.7-SC were tested for alcohol-induced LORR and alcohol drinking behavior. Similar to cuprizone mice, site-specific knockdown of *Ndrg1* within PFC confirmed a significant increase in acute high-dose alcohol sensitivity compared to scrambled control (Fig. 4A, *P =* 2.1 e-02). To examine alcohol sensitivity involved with drinking behavior, animals were tested using an intermittent three-bottle choice paradigm that included water, 15% alcohol, or 30% alcohol. Knockdown of *Ndrg1* significantly increased 15% alcohol consumption (Figure 4B, *P* < 1.0 e-04) while significantly decreasing 30% alcohol consumption (Figure 4C, *P* < 5.0 e-03). Animals self-titrated the overall amount of alcohol consumed in the three-bottle task, displaying no difference in total consumption between pLL3.7-*Ndrg1*-kd and pLL 3.7-SC. However, when 30% alcohol intake is analyzed as a proportion of 15% alcohol intake, *Ndrg1*-kd mice show a significant (Fig. 4D, *P* < 1.0 e-04) decrease in preference for 30% alcohol. In summary, to achieve comparable levels of total alcohol consumption *Ndrg1*-kd mice display a pattern of avoiding higher percent alcohol concentrations. Viewed alongside the effect of *Ndrg1*-kd on LORR, our findings reveal reduced PFC expression of *Ndrg1* is capable of modify sensitivity to higher alcohol concentrations, which may affect initial and long-term behavioral traits related to alcohol addiction.

**Figure 4:**
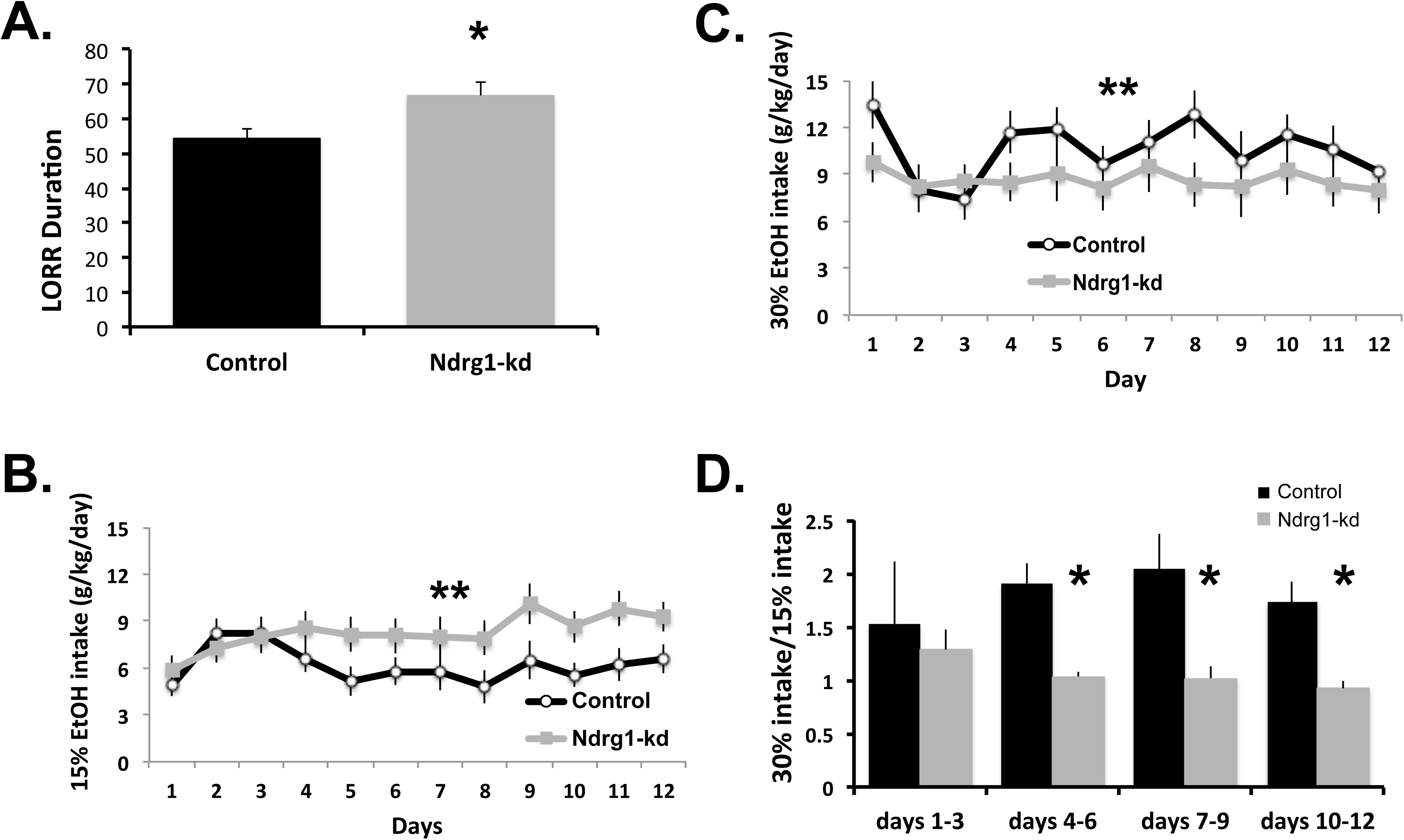
Alcohol behavioral effects of *Ndrg1*-knockdown. **(A)** Lenti-viral mediated knockdown of *Ndrg1* within the PFC increased the duration of alcohol-induced LORR (*P* = 2.1 e-02, n=16-20/group). *Ndrg1*-knockdown mice showed **(B)** significantly increased 15% ethanol consumption (*P* < 1.0 e-04, n=18-20/group), **(C)** significantly decreased 30% alcohol consumption (*P* < 5.0 e-03, n=18-20/group) **(D)** When 30% alcohol intake is analyzed as a proportion of 15% alcohol intake there is a significant decrease in the preference for 30% alcohol versues 15% (*P* < 1.0 e-04, n=18-20/group). These results within the PFC demonstrate a significant role for *Ndrg1* for alcohol behavioral sensitivity.

## Discussion

The PFC is an important brain-region involved in addiction to alcohol and substances of abuse (51), as well as several neuropsychiatric and neurodevelopmental disorders. Applying a systems genomics approach within PFC across humans and mice our analysis determined an evolutionary conserved gene coexpression network revolving around the prioritized candidate gene *Ndrg1*. The identified coexpression network was strongly enriched for multiple myelin-associated genes, including myelin basic protein (*Mbp*) and proteolipid protein (*Plp1*), which are both prominent structural constituents of the myelin sheath. Altering expression of myelin-associated genes *in vivo*, and individually targeting *Ndrg1* within PFC, showed a causal involvement for genes in alcohol sensitivity and alcohol consumption.

An inverse relationship between alcohol sensitivity and risk for development of AUD has been repeatedly observed in humans (52) (53), and may be an important factor in prevention based programs (54; 55). The neurobiology governing such behavior remains unclear; however, white matter (i.e. myelin) within PFC may be a critical neurobiological substrate. Functional magnetic resonance imaging (fMRI) studies have shown the PFC is less affected by alcohol exposure in individuals with a family history of alcoholism (56). Myelinated fiber integrity is increased within the frontal cortex of adolescents with a family history of alcoholism (57). Variation in myelin abundance can also affect subtle differences in cognitive and behavioral performance (14; 15; 58); suggesting neuronal-glia interactions involving myelin within PFC is a crucial component of CNS-mediated behaviors. Chronic alcohol abuse can severely affect myelin structure and function (7; 12; 59) that may partially recover during periods of prolonged abstinence from alcohol (60) (61). In the context of our currents studies this suggests dynamic alterations in the amount of myelin may be both, an important risk factor for the initiation of excessive alcohol consumption and a vital component of CNS plasticity underlying an AUD.

Despite the important role of myelin within CNS, the precise function of many myelin-related genes are still uncertain (62). Our studies further support the involvement of *Ndrg1* with myelin-associated gene expression, and suggest a novel role of *Ndrg1* as a determinant for alcohol sensitivity and intake. *Ndrg1* has an uncharacterized role in neuronal-glial cell differentiation and myelination (47) (63). Over stimulation of the proto-oncogene Myc inhibits *Ndrg1* expression, leading to progressive hypomyelination (64). NDRG1 is proposed to regulate cholesterol transport and associated intracellular trafficking mechanisms (65) (66). Myelin composition is fundamentally different from the cell membrane, enriched with differing myelin structural proteins, glycosphingolipids, and cholesterol (67). Through regulation of protein-lipid interactions and trafficking to sites of neuronal ensheathment *Ndrg1* may be a sensor controlling CNS myelin variation.

Myelin within PFC is an important facet of CNS-mediated executive function and motor control (68) (69). Inhibition of activity-dependent myelination causes decrements in motor learning paradigms (58). CNS myelin abundance is balanced by excitatory neuronal activity, which contributes to active myelination and proliferation of oligodendrocytes precursor cells. Optogenetic activation of neurons within adult mouse PFC increases the thickness of myelin sheaths and enhances motor function (70). Excitatory glutamatergic neurons can synapse with oligodendroglia cells, facilitating intracellular communication (71). PFC differences in baseline myelin-associated gene expression were also associated with glutamatergic-related systems in our current study. Additionally, the *Ndrg1* coexpression network was significantly enriched for genes expressed in glutamatergic neurons. *N*-methyl-*D*-aspartate (NMDA) glutamate receptors are known targets of alcohol action (72; 73), contributing to alcohol sensitivity and the development of alcohol dependence (74). Fyn kinase, a non-receptor tyrosine kinase that interacts with the NMDA-receptor, null mice are more sensitive to LORR (39) (75). In addition to affecting NMDA-receptor function, Fyn kinase plays an active role in CNS myelination with *Fyn* null mice having reduced myelin-associated gene expression (including *Ndrg1*) (76). Manipulating the expression of *Ndrg1* or myelin-related genes mirrors the LORR behavior witnessed in *Fyn* null mice. Variation in myelin may serve as a parallel mechanism to glutamatergic function, contributing to salient neuronal-glial cell interactions that help shape alcohol sensitivity and long-lasting drinking behavior.

Differences in the abundance of myelin-associated gene expression periods may confer vulnerability to a wide spectrum of psychiatric and neurological disorders (77–81). Commonalities involving myelin gene expression between individuals afflicted with an AUD and other psychiatric disorders suggest a comorbid pathophysiology related to oligodendrocytes and neuronal-glia dysfunction. Individual genes may participate in numerous cellular and behavioral systems. Our results highlight *Ndrg1* as a candidate gene affecting behavioral sensitivity to alcohol. Although additional evidence is necessary for uncovering cellular signaling mechanisms involving *Ndrg1* in the regulation of alcohol behavioral phenotypes, our results underscore the importance of *Ndrg1* and myelin-associated gene expression in the neurobiology of disease. Multiple myelin-related genes are correlated with *Ndrg1* across species, which may signify a quantitative trait gene network (45) within PFC influencing alcohol behavioral sensitivity. *Ndrg1* and closely related molecular systems (e.g. myelin) are critical components in the initial and chronic stages of alcohol use. Continued investigation of myelin will lead to a better understanding of diverse neurobiological disorders and potentially help ameliorate disease.

## Acknowledgements

This research was supported by the National Institutes of Health (NIH) – National Institute on Alcohol Abuse and Alcoholism (NIAAA): U01AA016667, P50AA022537, and R01AA020634 TO MFM. The authors would like to thank Drs. Beth Bennett, Tom Johnson and Chris Downing from the Institute of Behavioral Genetics, University of Colorado Bolder for assistance in formulating preliminary studies on ILS and ISS mice. Additionally, Dr. Keith Shelton from the Department of Pharmacology and Toxicology at Virginia Commonwealth University provided technical assistance for blood ethanol determinations.

## Supplementary Figure Legends

**Supplementary Figure 1: Independent assessment of genetic differences between ILS and ISS related to *Ndrg1* expression.** Analysis of *Ndrg1* basal gene expression across the LXS RI panel from **(A)** whole brain (n = 62) and **(A)** hippocampus (n = 79); showing a significant cis-eQTL for *Ndrg1* in this mouse population. In general, the larger the number of animals available for study increases the ability to detect a cis-eQTL. **(C)** Comparison of the ISS and ILS mouse genome shows *Ndrg1* resides in a non-identical by descent region (white) within the LORR behavioral QTL region of chromosome 15. ISS and ILS genetic data is from the mouse phylogeny viewer (http://msub.csbio.unc.edu/) (85), with regions identical by descent (IBD) between ISS and ILS shown in pink.

**Supplementary Figure 2:** The mean alcohol-induced duration of LORR across A/J, C57BL/6J, FVB/NJ, DBA/2J, 129S1/SvlmJ, BALB/cByJ, and C3H/HeJ mice (n=5-6/strain) in our current study is significantly correlated (Pearson’s r = 0.91, *P* = 4.8 e-03) with a prior report of alcohol-induced LORR conducted in these strains of mice (46).

**Supplementary Figure 3: Cuprizone does not affect taste preference for sweet or bitter solutions. (A)** Saccharin intake and **(B)** quinine intake were not significantly different over five days of 2-bottle choice for water or tastant. The average intake of **(C)** saccharin (*P* = 3.6 e-01, n=8/group) or **(D)** quinine (*P* = 3.23 e-01, n=8/group) were not different between cuprizone and controls. These data suggest that cuprizone does not significantly alter taste preference.

**Supplementary Figure 4:** Blood ethanol levels in control and cuprizone mice are not significantly different over 4 hours (main effect of time, *P* < 1.0 E-04, n=4/group/time). These results indicate cuprizone does not have off-target effects on the metabolism of alcohol.

**Supplementary Figure 5: Cuprizone does not alter locomotor behavior or fine motor coordination.** Cuprizone does not alter basal or alcohol-induced locomotor activity as measured by **(A)** distance traveled, **(B)** average velocity or **(B)** ambulatory episodes following 2 g/kg (i.p.) alcohol or saline (n=14/group). Main effect of alcohol administration on average velocity (*P* < 1.0 e-04), but there are no significant interactions between the groups. **(D)** Gait analysis was conducted to determine potential effects on fine-motor coordination (n=10/group). No effect of cuprizone treatment was noted for **(E)** stride length (*P* = 7.53 e-01) 0.753), **(F)** stance length (*P* = 9.24 e-01) or **(G)** sway length (*P* = 8.73 e-01), suggesting there are no adverse effects of cuprizone on mouse locomotor or gaiting behavior.

**Supplementary Figure 6: Cuprizone effect on handling induced convulsion behavior.** Blinded observation of HIC activity revealed cuprizone had no effect on **(A)** baseline HIC (*P* < 5.33 e-01) or **(B)** alcohol-withdrawal HIC time-course (main effect of time, *P* < 1.0 e-04). These results show that cuprizone induced demylination has no adverse generalized CNS hyperexcitability. Data Mean ± SE.

**Supplementary Figure 7: Confirmation of Ndrg1 knockdown. (A)** In NIH/3T3 cells, *Ndrg1* mRNA was significantly decreased in pLL3.7-Ndrg1-kd compared to pLL3.7-SC as measured by qRT-PCR (*P* = 1.8 e-03, n=3/group). **(B)** pLL3.7-Ndrg1-kd lentivirus significantly decreased NDRG1 protein levels compared to the pLL3.7-sh-SC virus by western blot normalized to β–actin (*P* < 1.0 e-04, n=3/group). **(C)** Stereotaxic injection of 7 x 10^-6^ TU/μl pLL3.7-SC or -Ndrg1-kd lentivirus into the PFC of C57BL/6J mice (coordinates 0.6 mm lateral, 1 mm anterior and 2 mm ventral to bregma). Mice were sacrificed 3 weeks after injection to verify the site of injection and NDRG1 knockdown. eGFP expression (green) and NDRG1 protein (red) knockdown via immunohistochemistry.

## Supplementary Table Legends

**(Supplementary Tables available from authors on request)**

**Supplementary Table 1:** Complete list of basal differences in gene expression within the PFC for ISS versus ILS mice and B6 versus D2 (6). The gene lists have also been publically shared through http://geneweaver.org/ (25); however, please be aware that there are small differences in lists due to gene annotation. These differences are minor and do not change the reported results and conclusions.

**Supplementary Table 2:** Gene ontology analysis of consistent basal differences in gene expression within the PFC (Fig. 1A, Supplementary Table 1); showing genes related to the predisposition for differences in alcohol behavioral sensitivity are involved in diverse biological categories, notably member of the ‘myelin sheath’ and components of ‘glutamate receptor signaling.’

**Supplementary Table 3:** Aggregate of genes significantly correlated (*P* < 0.05) with *Ndrg1* gene expression within the PFC across the BXD RI panel (n=29), LXS RI panel (n=42), and humans (n=31). The *Ndrg1* gene coexpression network is enriched for multiple myelin-related genes. Gene tested using qRT-PCR following a cuprizone diet (Fig. 2) are in **bold.** The gene list is ordered by the cumulative rank of strongest correlation across all three datasets.

**Supplementary Table 4:** Over-representation of genes within the PFC that are mutually correlated to *Ndrg1* LXS and BXD phenotypes that possess a sufficient number of mutual strains of mice (n ≥ 3). BXD mice are in general more widely studied in relation to LXS mice, respectively providing 4,372 and 189 phenotypes for our analysis. The phenotypes for both RI panels are publically available at http://genenetwork.org/webqtl/main.py. Each phenotype is assigned a unique GeneNetwork ID within the respective RI panel, and if available corresponds to a published report. Not all phenotypes were significantly related to the *Ndrg1* coexpression network; however, the gene network was strongly related to LORR in both the BXD and LXS RI panel, as well as alcohol-related drinking behavior across species.

**Supplementary Table 5:** Summary of alcohol behavioral effects for animals continuously exposed to cuprizone, an animal model of CNS demyelination.

