## Supplementary figures and images for "Variation of Myelin-associated Gene Expression and *Ndrg1* within the Prefrontal Cortex as Determinants for Initial Level of Response to Alcohol"

### Supplementary Fig. 1

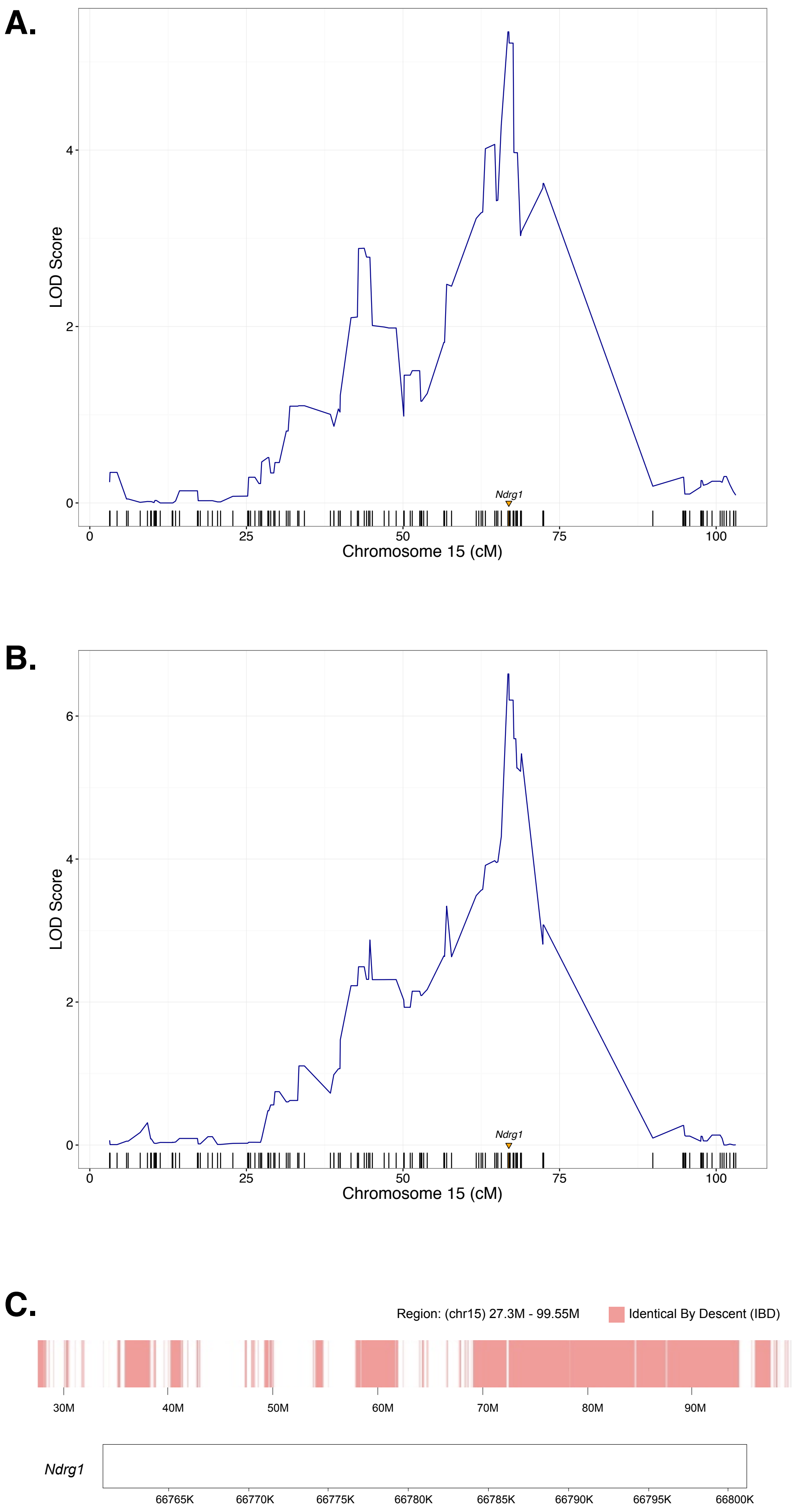

### Supplementary Fig. 2

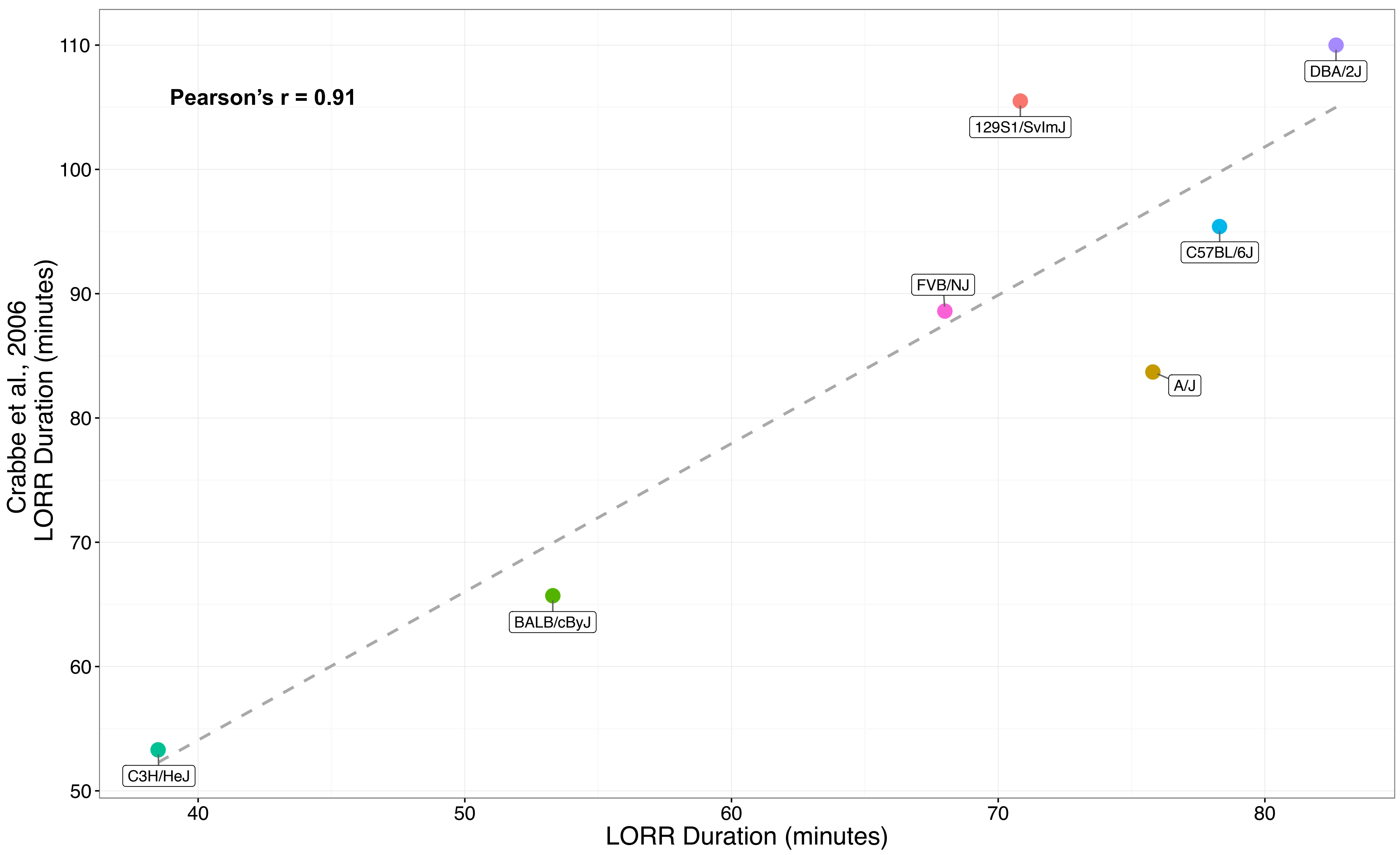

### Supplementary Fig. 3

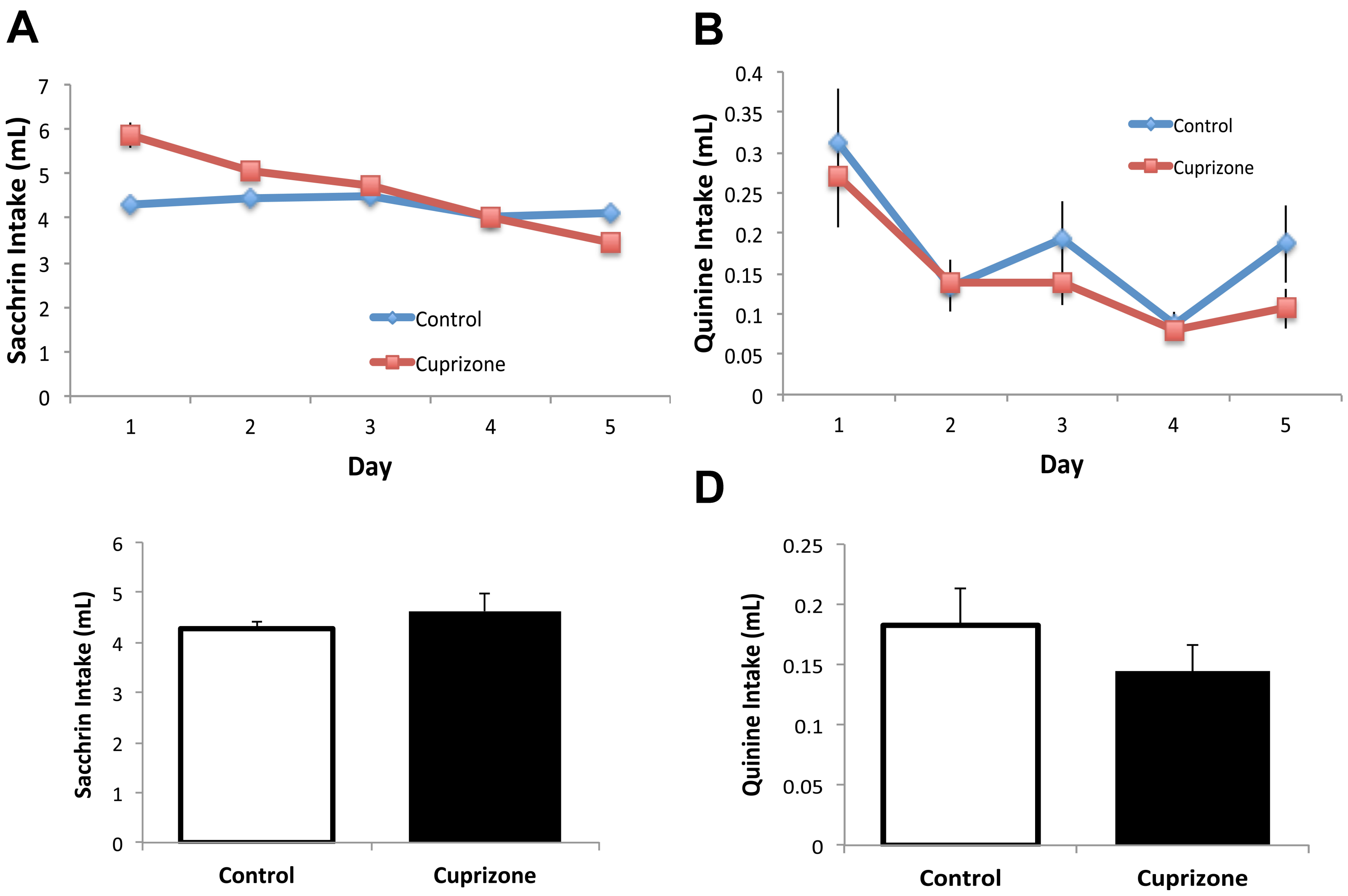

### Supplementary FIg. 4

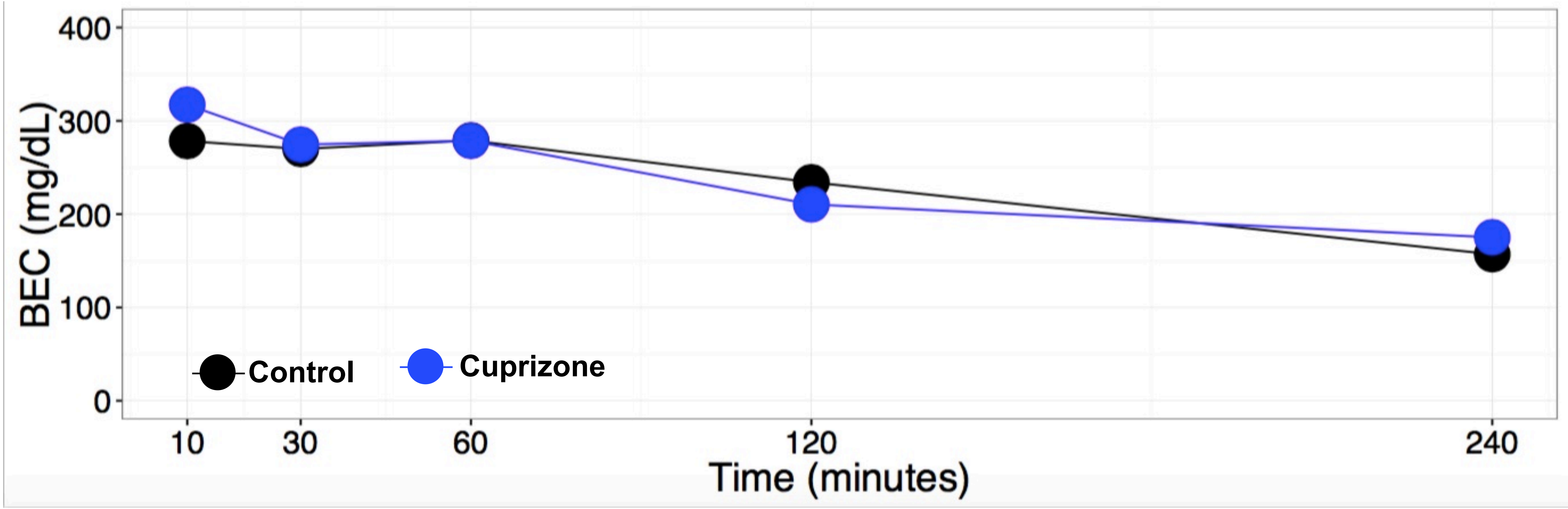

### Supplementary Fig. 5

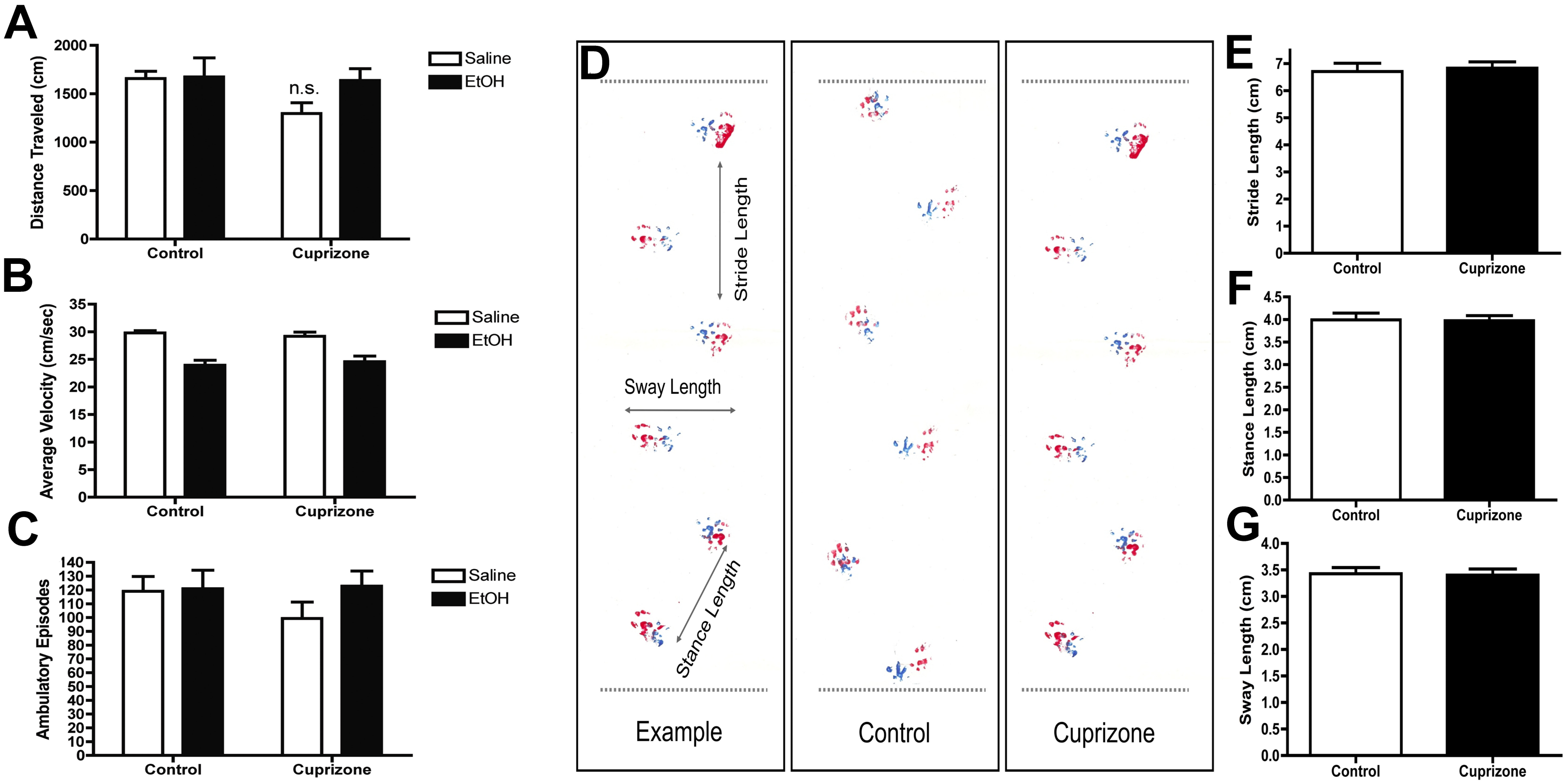

### Supplementary FIg. 6

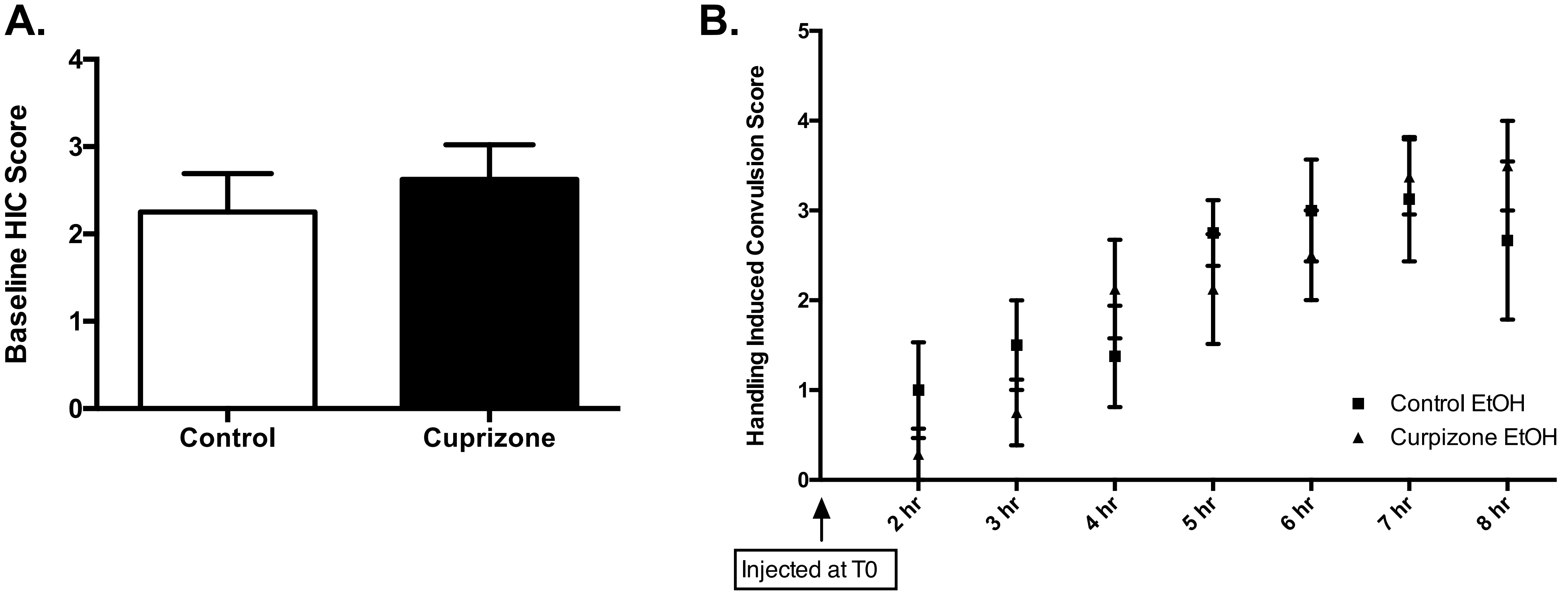

### Supplementary Fig. 7

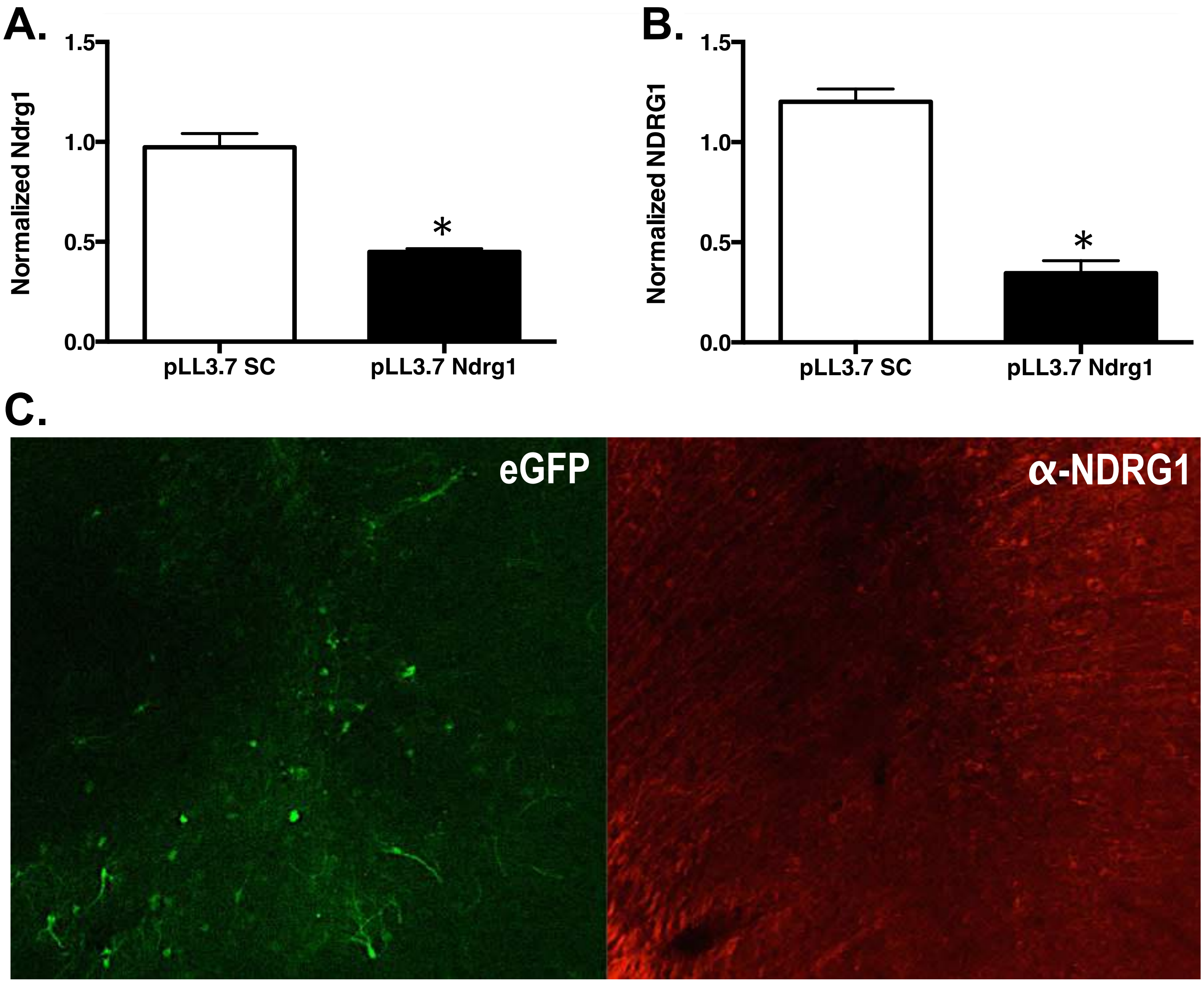
