## Supplemental Methods for "Variation of Myelin-associated Gene Expression and *Ndrg1* within the Prefrontal Cortex as Determinants for Initial Level of Response to Alcohol"

**Quantitative real-time reverse transcription – qPCR:** Total RNA (1 µg) was reverse transcribed into cDNA using iScript cDNA synthesis kit (Bio-Rad). Quantitative real-time PCR performed using iQ SYBR Green Supermix and the CFX system (Bio-Rad) according to manufacturer's instructions. Three technical replicates were performed for each biological sample of cDNA, with therelative abundance of target transcripts normalized to the housekeeping gene *Gapdh.* Primers were designed to minimize secondary structure formation and cross intron-exon boundaries to reduce genomic DNA amplification.

**Protein expression of NDRG1:** Cells were lysed in RIPA buffer supplemented with HALT Protease Inhibitors (Fisher Scientific). Protein concentrations were determined using BCA Protein Assay (Pierce). Equal amounts of protein were loaded per lane and separated on a 10% SDS–PAGE gel. Protein was transferred to a PVDF membrane. We detected NDRG1 using antibody against NDRG1 (AbCam #Ab63989) and donkey antibody against rabbit conjugated with horseradish peroxidase. The blot was developed with ECL+ reagent (Amersham Biosciences) and visualized on XoMAT film. NDRG1 protein expression was normalized to β-actin (A2228, Sigma). Immunohistochemistry for NDRG1 was performed in mice perfused with 4% paraformaldehyde as previously described (Costin 2013). Microscopy was performed at the VCU Department of Anatomy and Neurobiology Microscopy Facility, supported with funding from the NIH-NINDS Center core grant (5P30NS047463).

**Fluoromyelin staining:** After 5 weeks on cuprizone diet or normal chow, mice were perfused with 4% paraformaldehyde, cryoprotected in sucrose and frozen. Brains were sectioned (30 □m) on a cryostat and stained with fluoromyelin (Life Technologies) according to the manufacturers protocol.

**Loss of righting reflex (LORR) due to alcohol**: Following 3 days of habituation to intraperitoneal (i.p.) 0.9 N saline injections, mice were administered a 4 g/kg dose of alcoho (i.p.) using a 20% (v/v) stock solution and placed in a clean cage with bedding. Each mouse was timed for the onset of the LORR from the moment of injection until acquiring the LORR. Any animals not acquiring LORR within two minutes were deemed an outlier and removed from the study. Animals were deemed to have acquired the LORR when placed on their back V-shaped cage-top and remained in that position for no less than 20 seconds. Following acquiring the LORR animals were scored for the duration of the LORR, defined as the amount of time it takes an animal to right itself onto all four paws 3 times within 60 seconds.

**2-bottle choice alcohol consumption and taste perception:** Mice were individually housed and given access to two bottles in their home cage. 10% (w/v) alcohol was given at 5 weeks for 15 days, followed by 20% (w/v) for 13 consecutive days. After the completed collection of drinking data, taste perception test was done with the same 2-bottle choice paradigm for 2mM saccharine or 126 µM quinine for 5 consecutive days.

**Intermittent 3-bottle choice alcohol consumption:** Mice were individually housed with four weeks of intermittent access to 15% and 30% alcohol solutions supplementing a water bottle. Alcohol was placed in cages on Mondays, Wednesdays, and Fridays at the beginning of the 12-h dark cycle, with measurements of alcohol intake taken on Tuesdays, Thursdays, and Saturdays. Total amount of each alcohol solution and water were measured and alcohol intake and preference were calculated within each group of mice.

**Locomotor behavior:** Animals habituated to the behavioral room for one hour prior to testing and testing was conducted between 8 AM and 12 PM EST. Mice were given saline or 2 g/kg alcohol i.p. and placed immediately inside the behavioral chamber. Behavioral boxes were enclosed in a sound-attenuating chamber equipped with overhead lighting in and ventilation system, interfaced with Med Associates software. Beam breaks were tested for 10 minutes in the chamber using 5 minute bins for each group of animals.

**Gait analysis of mouse locomotor behavior:** Ataxia and fine-motor coordination were determined using the footprint test, also referred to as gait analysis, (Crawley, 1999) for mice on normal rodent chow and food supplemented with cuprizone. Pronounced loss of CNS myelin can impair fine-motor control in mice (Jensen et al., 1993, Emery et al., 2009). Following habituation to the testing room, mice treated +/- cuprizone in standard chow had their paws painted with non-toxic paint. Front paws were painted with blue and rear paws were painted in red. On day one, each animal was immediately placed the gait analysis box and allowed to run to the end of the chamber. Following this initial run animals were each given a second opportunity to run through the chamber, returned to their home cage, then given a i.p. saline injection and run the third time representing time zero. Each animal was run again at 5 minutes, 15 minutes, 30 minutes, and 60 minutes under saline conditions to complete day one. On day two for each group each animal was given an i.p. 2 g/kg ethanol injection and immediately placed in the chamber for time zero then run again at 5 minutes, 15 minutes, 30 minutes, and 60 minutes. On day three for each group each animal was given an i.p. 2.75 g/kg ethanol injection and immediately placed in the chamber for time zero then run again at 5 minutes, 15 minutes, 30 minutes, and 60 minutes. Animal groups were rotated to give each group at least one day off prior to repeated testing. Paper strips for each animal were hung to dry and measured using a standard ruler for stride length, stance length, and sway length. Measurements were sampled using the average of six repetitive steps from the midsection of the paper.

**Lentivirus generation and characterization:** Commercially available plasmids for shRNA to induce knockdown of *Ndrg1* and a scrambled control vector were purchased commercially (NDRG1 MISSION shRNA Bacterial Glycerol Stock, SHC002 and MISSION pLKO.1-puro Non-Mammalian shRNA Control, SHC001, Sigma Aldrich). The sense and scrambled oligomers for *Ndrg1* and the human U6 promoter were cut out of the pLKO.1 vector and cloned into the pLentiLox 3.7(Addgene, plasmid 11795) vector using XhoI and XbaI. The human U6 promoter was retained from the pKLO.1 vector since it has higher infection efficiency than the mouse U6 promoter (Roelz 2010). The pLL3.7 vector was selected since it had an eGFP site to allow localization. Ligation products were transformed into competent E. coli DH5a cells, then cultured in LB agar plates to select for colonies with inserted shRNA sequences using colony PCR. Proper insertion into the plasmid was verified by sequencing the plasmid from the human U6 promoter. Replication-defective lentiviruses were generated by transient co-transfection of 293FT cells with three plasmids (pLL3.7-SC or Ndrg1-kd, psPAX2-packaging plasmid, pMD2.G-VSV-G envelope expressing plasmid, Addgene, Cambridge, MA) using Lipofectamin 2000 mediated transfection. Mouse fibroblast (NIH3T3) cells were then infected with virus and knockdown of *Ndrg1* was verified by qPCR and western blot, normalized to *Ppp2r2a* and *Nduvf* as reference genes and β-actin as reference protein.
